# Integrating Fas-mediated apoptosis with IFNγ signaling to drive tumor regression in mRNA cancer therapeutics

**DOI:** 10.64898/2026.04.06.716844

**Authors:** Hee-Su Shin, Soon-Gyu Kwon, Hayyoung Lee, Jie-Oh Lee

## Abstract

For mRNA-based cancer gene therapy, we engineered a membrane-bound fusion protein combining interferon-γ (IFNγ) with the Fas intracellular domain (FasICD) to couple local IFNγ signaling with Fas-driven apoptotic tumor cell death. IFNγ–FasICD was robustly expressed on the plasma membrane after mRNA transfection. In murine cancer cell lines, IFNγ–FasICD mRNA reduced viability within 24 h, resulting in ∼50% cell death in MC38 cells and ∼75% in B16OVA cells, exceeding the cytotoxicity of the FasICD-deleted control (IFNγ–FasΔ). Mechanistically, IFNγ–FasICD induced predominantly apoptotic rather than necrotic cell death. IFNγ–FasICD also activated IFNγ receptor signaling in both cancer and the immune cells, inducing IFNγ-responsive genes in IFNγR-high B16OVA cells and triggering STAT1 phosphorylation in co-cultured splenocytes. For *in vivo* delivery, IFNγ–FasICD mRNA was formulated in lipid nanoparticles (LNPs), enabling strong intratumoral expression that peaked at ∼3 h and persisted for more than 48 h. Repeated intratumoral injections of LNP-formulated IFNγ–FasICD mRNA suppressed the growth of established B16OVA and MC38 tumors and improved survival, with ∼40% and ∼20% of mice surviving beyond 30 days, respectively. IFNγ–FasICD treatment remodeled the tumor microenvironment by increasing tumor-infiltrating CD45^+^ cells and CD8^+^ T cells, while further reducing FOXP3^+^ regulatory T cells. Moreover, NK/NKT cells and cDC1/cDC2 populations were increased, and their activation was enhanced. In tumor-draining lymph nodes, IFNγ–FasICD mRNA promoted dendritic cell migration and increased priming and differentiation of CD8^+^ T cells toward effector and memory phenotypes, accompanied by enhanced functional activation of IFNγ-producing CD8^+^ T cells and highly cytotoxic NK cells in peripheral blood. Overall, our findings provide a mechanistic foundation for cytokine–death receptor fusion proteins as an *in vivo* antitumor strategy that can reprogram tumor cells into localized sources of both apoptotic signals and immune-activating cues.

## Introduction

Gene therapy for cancer involves introducing modified genes to treat cancer by repairing a faulty one or enhancing the immune system to target cancer cells (1–3). The strategy of directly targeting cancer cells has the advantage of inhibiting cancer cell proliferation and inducing apoptosis, thereby minimizing damage to normal cells, a major limitation of conventional chemotherapies and radiotherapies. In addition, it can induce immune responses against a broad range of tumor antigens released from dying cancer cells, which is advantageous for overcoming tumor heterogeneity (4–6). Conventional DNA-based or oncolytic viruses use DNA vectors to deliver genetic material to cells (7, 8). On the other hand, mRNA cancer therapy uses mRNA to instruct the immune system to fight cancer, primarily through vaccines that present tumor proteins as antigens, or by engineering immune cells to recognize and kill cancer, offering a personalized and powerful way to activate the body’s own defenses (9, 10). mRNA-based gene therapy has the advantages of lacking the risk of genomic modification, enabling rapid protein expression, and allowing easy mass production through *in vitro* transcription processes (11, 12). In addition, it can be stably delivered *in vivo* based on LNPs and can be applied to tissue-specific targeting or APC targeting (13–15). Based on this, mRNA technology has been efficiently used to express cancer vaccines encoding tumor-specific antigens or cytokines.

Pro-apoptotic gene delivery can be performed or an oncolytic virus can be used as a method to induce apoptosis in cancer cells, and the anti-apoptotic gene of cancer cells can be suppressed based on siRNA/shRNA to promote apoptosis (16–18). Among them, the pro-apoptotic gene delivery method uses the death receptor/ligand pathway such as FasL, TRAIL receptor, and TNF alpha or overexpresses p53, BCL-2 family gene, caspase-based genes (19–21). It is possible to target only cancer cells by applying the recently advanced *in vivo* gene delivery technology, especially with the tumor specific targeted nanobody or liposome to which the ligand of the cancer cell-specific receptor is attached (22). Therefore, effective anticancer effects can be expected while minimizing normal cell damage. Fas (CD95) is one of the representative cell death receptors belonging to the TNF receptor superfamily, and when combined with Fas ligand (FasL, CD178), an intracellular domain, the fas-associated death domain (FADD) binds to the death domain, causing apoptosis through caspase after DISC formation (23). Cancer cells can be resistant to FasL because of the low expression of Fas in cancer (24). Then it can be overcome by overexpressing Fas specifically or by overexpressing Fas-mediated chimeric protein (25–27).

IFNγ is mainly secreted by activated T cells and NK cells and plays a role in enhancing the antitumor immune responses (28, 29). IFNγ is involved in M1 polarization and induces IL-12 production by macrophages and dendritic cells to form a positive feedback loop that stimulates T cells and NK cells again (30, 31). IFNγ acts as a potent antitumor cytokine by directly inducing apoptosis and cell cycle arrest in various cancer cells, typically mediated through the JAK1/2-STAT1 signaling pathway (32, 33). In addition, by increasing the expression of MHC class I and class II molecules, it is possible to improve tumor antigen presentation of cancer cells (34, 35). However, since IFNγ receptors are not only expressed in almost all immune cells but also in fibroblasts, epithelial cells, and endothelial cells, chronic and systemic IFNγ exposure can lead to serious side effects (36–38). To overcome this, various methods of delivering IFNγ are being developed to limit their expression in tumor microenvironment (39–41). Among them, the method of expressing cytokines in a membrane-bound form has the advantage of stably expressing cytokines in the tumor microenvironment (42–46).

In our previous studies, we showed that IFNγ secreted by IL-9–induced macrophages promotes immune-cell recruitment into the tumor microenvironment (TME) and drives M1 macrophage polarization, thereby enhancing antitumor activity (47). We also reported that CT26 tumor cells engineered to stably express a membrane-bound IFNγ–Fas fusion protein functioned as an effective cancer vaccine: compared with IFNγ alone (48). However, cell-based vaccine strategies face practical limitations for broad therapeutic use, including complex manufacturing, quality control, and scalability (49). In the present study, we substantially advance this concept by converting the IFNγ–Fas fusion into an LNP-formulated mRNA therapeutic for intratumoral administration to established solid tumors. This platform enables rapid, transient, and localized expression of membrane-bound IFNγ–FasICD directly in tumor cells, effectively reprogramming the tumor into an in situ source of both immune-stimulatory cytokine signaling and pro-apoptotic death signaling. *In vivo*, intratumoral administration of IFNγ–FasICD mRNA-LNP suppressed tumor growth and improved survival, accompanied by consistent remodeling of the TME, including increased infiltration and activation of immune effector populations and a reduction in FOXP3^+^ regulatory T cells. Moreover, treatment facilitated dendritic cell trafficking and a bias toward effector and memory differentiation within tumor-draining lymph nodes, while increasing the effector potential of circulating CD8^+^ T cells and NK cells. Collectively, the cytokine–death receptor fusion proteins delivered as mRNA-LNPs represent an *in vivo* antitumor strategy that integrates direct tumor cell apoptosis with coordinated immune activation, while overcoming key feasibility barriers associated with cell-based cancer vaccines.

## Results

### Expression of IFNγ and the Fas intracellular domain fusion protein induces apoptosis in murine cancer cells

IFNγ and the Fas intracellular domain (FasICD) were fused via the Fas transmembrane domain to enable membrane localization of the fusion protein (Figures 1A, B). IFNγ-FasΔ, in which the FasICD is deleted, was used as a control to assess Fas-mediated apoptotic effects *in vitro*. HEK293T cells were transfected with mRNA encoding either IFNγ-FasΔ or IFNγ-FasICD, and IFNγ expression levels were evaluated in whole-cell lysates and on the cell surface by ELISA and flow cytometry, respectively. IFNγ was robustly expressed in IFNγ-FasICD mRNA–transfected cells, both in total lysates and on the plasma membrane (Figures 1C, D). IFNγ was also detected in the culture supernatant, indicating partial proteolytic processing of the fusion protein and release of soluble IFNγ (Figure 1E). Because membrane-bound IFNγ expression was clearly confirmed and the additional soluble IFNγ may further enhance antitumor immune activity in the tumor microenvironment, subsequent experiments were performed using the IFNγ-FasICD fusion construct.

**Figure 1.**
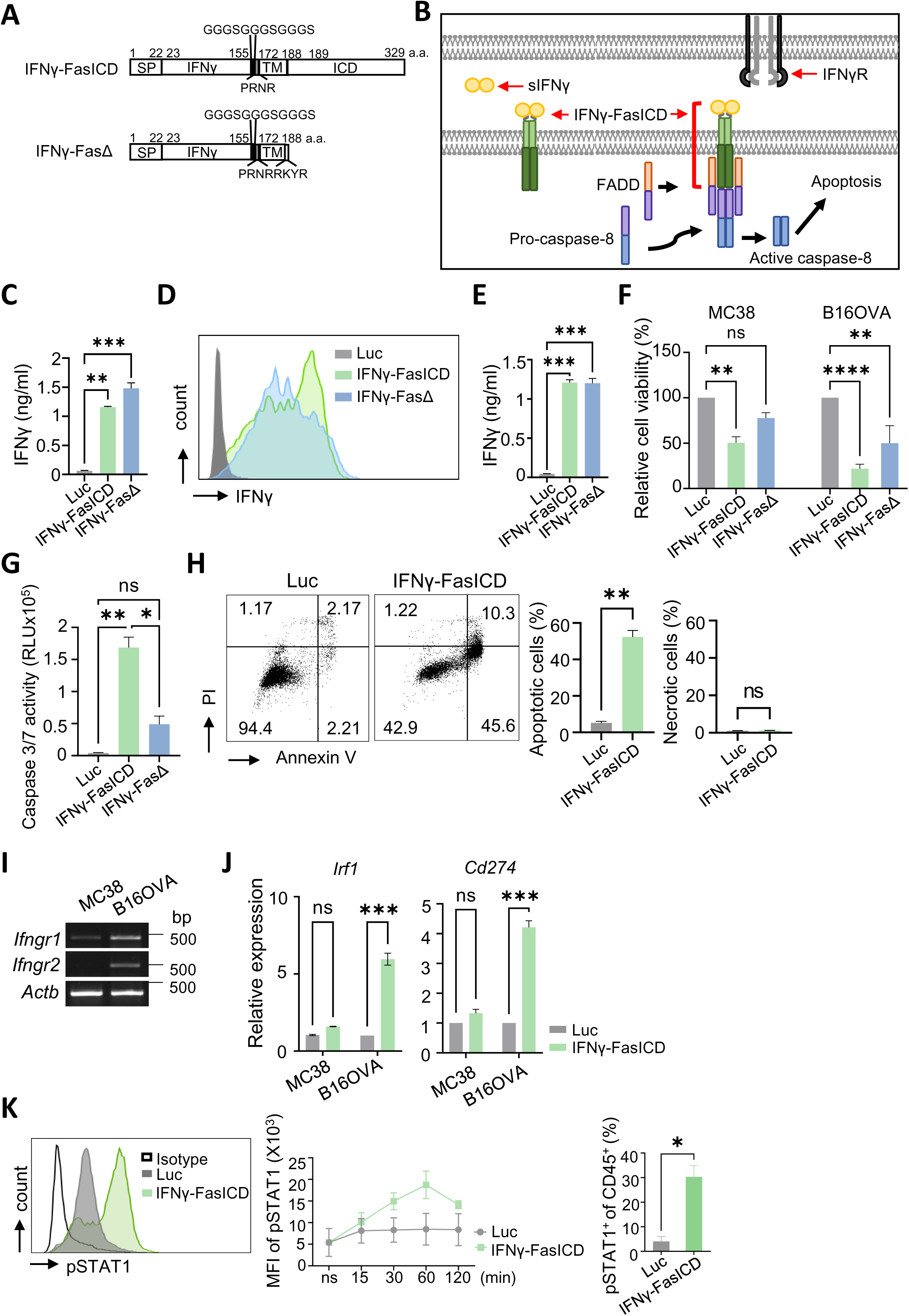
Expression and functional validation of a membrane-bound IFNγ–FasICD fusion protein inducing STAT1 phosphorylation and Fas-mediated apoptosis. (**A**) Schematic diagram of IFNγ-FasICD and IFNγ-FasΔ proteins. SP; The IFNγ signal sequence peptide, FasTM; Fas transmembrane domain, FasICD; Fas intracellular domain. A linker (GGGSGGGSGGGS) was added. Numbers indicate amino acid positions. **(B)** Model of IFNγ-FasICD working mechanism. (**C-E**) HEK293T cells were transfected with IFNγ-FasICD mRNA. **(C)** IFNγ levels in cell lysates were measured by ELISA at 6 h. **(D)** The expression of membrane-bound IFNγ was analyzed using flow cytometry. **(E)** IFNγ levels in culture supernatants were measured at 6 h. **(F)** MC38 and B16OVA cells were transfected with Luc (luciferase), IFNγ-FasICD, or IFNγ-FasΔ mRNA, cell viability was assessed by MTT assay at 24 h. (G-H) B16OVA cells were transfected with the IFNγ-FasICD mRNA. Caspase-3/7 activity was measured to assess apoptosis **(G)** and Annexin V^+^ cells were quantified **(H)**. **(I)** Expression levels of IFNγ receptor 1 and 2 (*Ifngr1* and *Ifngr2*) were analyzed by semi-quantitative RT-PCR. β-actin (*Actb*) was used as an internal control. (J) Expression of IFNγ downstream target genes, *Irf1* and *Cd274*, was analyzed by real-time PCR. **(K)** B16OVA cells were transfected with IFNγ-FasICD mRNA and co-cultured with splenocytes from C57BL/6J mice. STAT1 phosphorylation (Tyr701) in splenocytes as analyzed over time by flow cytometry (left and middle panels), and the percentage of pSTAT1^+^ cells at the 60-min time point among CD45^+^ cells is shown (right panel). Data are pooled from three independent experiments. Statistical significance was determined by one-way ANOVA followed by Tukey’s multiple comparisons test (C, E, G), ordinary one-way ANOVA followed by Šídák’s multiple comparisons test (J), two-way ANOVA followed by Dunnett’s multiple comparisons test (F), and unpaired t-tests (H, K). Error bars indicate SEM. Note: \**P* < 0.05, \*\**P* < 0.01, \*\*\**P* < 0.001, \*\*\*\**P* < 0.0001.

MC38 colon carcinoma cells and B16OVA melanoma cells were first transfected with IFNγ–FasICD mRNA. Cell viability was assessed using an MTT assay. After 24 h, approximately 50% and 75% cell death was observed in IFNγ–FasICD mRNA–transfected MC38 and B16OVA cells, respectively (Figure 1F). Although IFNγ can support cell survival, it is also well established that it can induce tumor cell death and inhibit cancer cell growth. Consistent with this, transfection with IFNγ–FasΔ mRNA also induced approximately 25% and 50% cell death in MC38 and B16OVA cells, respectively, demonstrating IFNγ-mediated cytotoxicity in these cancer cell lines (Figure 1F). To further dissect the mechanism of cell death, caspase activity was measured following transfection with IFNγ–FasICD or IFNγ–FasΔ mRNA. Caspase activity was increased by approximately 3.5-fold in IFNγ–FasICD–transfected cells compared with IFNγ–FasΔ–transfected cells, indicating that FasICD strongly promotes apoptotic signaling in cancer cells (Figure 1G). These data suggest that although IFNγ alone can induce cancer cell death, a substantial fraction of this effect occurs through caspase-independent pathways, whereas FasICD predominantly triggers caspase-dependent apoptosis. Because necrosis within the tumor microenvironment is associated with poor prognosis and chronic inflammation and often promotes tumor progression through angiogenesis and cancer cell proliferation (50), we next examined whether IFNγ–FasICD–induced cell death occurred via apoptosis or necrosis. Annexin V/propidium iodide (PI) staining revealed that the majority of cell death induced by IFNγ–FasICD was apoptotic, with negligible necrotic cell death (Figure 1H). These results demonstrate that expression of the IFNγ–FasICD fusion protein induces predominantly apoptotic cell death in the transfected cancer cells.

### Expression of IFNγ-FasICD induces IFNγ receptor signaling in both murine cancer and immune cells

To determine whether membrane-bound or soluble IFNγ expressed by transfected cancer cells could activate IFNγ receptor signaling in cancer cells, IFNγ–FasICD–transfected B16OVA cells were analyzed for IFNγ downstream factors, *Irf1* and *Cd274* (encoding PD-L1). B16OVA cells exhibited relatively higher expression of IFNγ receptors (*Ifngr1* and *Ifngr2*) compared with MC38 cells (Figure 1I). The expression of IRF1 and CD274 was significantly induced in B16OVA cells by transfection with IFNγ–FasICD, whereas they were not induced in MC38 cells (Figure 1J). Then, IFNγ–FasICD–transfected B16OVA cells were co-cultured with splenocytes isolated from C57BL/6 mice to determine whether IFNγ expressed by transfected cancer cells could activate IFNγ receptor signaling in immune cells. Phosphorylation of STAT1 (pSTAT1, Tyr701), a major downstream effector of IFNγ receptor signaling, was assessed in the harvested splenocytes. Compared with splenocytes co-cultured with luciferase (Luc)-transfected B16OVA cells, co-culture with IFNγ–FasICD–transfected B16OVA cells induced STAT1 phosphorylation in a time-dependent manner, reaching a maximum of approximately 30% pSTAT1-positive cells among the CD45^+^ population (Figure 1K). These results demonstrate that expression of the IFNγ–FasICD fusion protein effectively activates IFNγ receptor signals in both autocrine and paracrine manners.

### Intratumoral injection of lipid nanoparticle (LNP)–encapsulated mRNA enabled efficient expression of IFNγ–FasICD in subcutaneously inoculated murine tumors

The LNP formulation consisted of the ionizable lipid SM-102, DSPC, cholesterol, and DMG-PEG2000 at a molar ratio of 50:10:38.5:1.5. The mRNA-incorporating LNP was characterized by dynamic right scattering (DLS), showing an average diameter of 101.7 nm with acceptable zeta potential and polydispersity index (Supplementary Figure 1A). The uniform size of LNPs was further confirmed by cryo-electron microscopy images (Supplementary Figure 1B). The encapsulation efficiency exceeded 95% (Supplementary Figure 1C). For intratumoral delivery, LNP-encapsulated mRNA was evaluated for *in vivo* expression in subcutaneous B16 tumors and compared with naked mRNA formulated in PBS or Ringer’s lactate solution (RLS). Following a single intratumoral administration of the LNP-encapsulated luciferase mRNA, bioluminescence imaging revealed peak signal intensity at 3 h post-injection, which remained robust for more than 48 h (Figures 2A, B). In contrast, no detectable bioluminescence signal was observed between 3 and 48 h after injection of naked mRNA formulated in either PBS or RLS. Similar kinetic analyses were performed in B16OVA–bearing mice following intratumoral injection of LNP-encapsulated IFNγ–FasICD mRNA. IFNγ–FasICD mRNA was administered either once and analyzed after 6 h (1X, 6 h), or twice at a 48 h interval and analyzed after 6 h (2X, 6 h) or 24 h (2X, 24 h) (Figure 2C). Following the experimental schedule outlined in Figure 2C, flow cytometric analysis of membrane-bound IFNγ expression on CD45^-^ tumor cells showed that approximately 10% of tumor cells were IFNγ^+^ across treatment conditions (Figures 2D, E). Because partial proteolytic processing of the fusion protein and subsequent release of soluble IFNγ can occur, the total number of IFNγ-expressing cells is likely underestimated and may be substantially higher.

**Figure 2.**
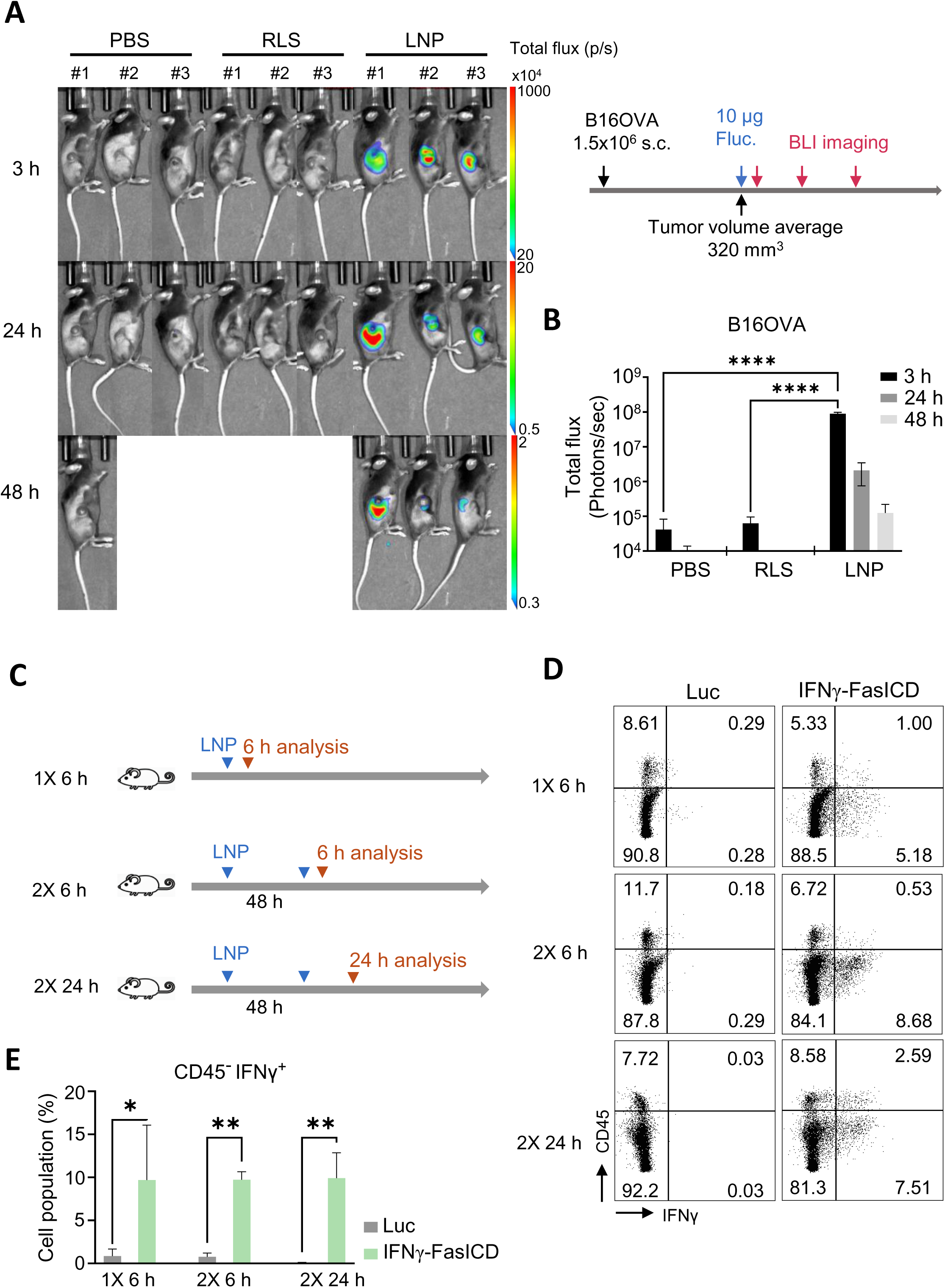
Intratumoral delivery of LNP-encapsulated mRNA enables efficient expression in subcutaneous murine tumors. **(A)** Representative images and a bar graph showing *in vivo* bioluminescence imaging (BLI) at 3, 24, and 48 h after a single intratumoral injection of luciferase mRNA in B16 tumors. mRNA was administered as naked mRNA in PBS, naked mRNA in Ringer’s lactate solution (RLS), or LNP-formulated mRNA (LNP) (*n* = 3). **(B–D)** Membrane-bound IFNγ expression in B16OVA tumors following intratumoral injection of IFNγ–FasICD mRNA was analyzed by flow cytometry (*n* = 3). **(B)** Experimental scheme. IFNγ–FasICD mRNA was administered once and analyzed 6 h later (1×, 6 h), or administered twice at a 48-h interval and analyzed 6 h (2×, 6 h) or 24 h (2×, 24 h) after the second injection. **(C)** Representative flow cytometry analysis of membrane-bound IFNγ expression. **(D)** Quantification of CD45^-^ IFNγ^+^ cells. Statistical significance was determined by two-way ANOVA followed by Šídák’s multiple comparisons test. Error bars indicate SEM. Note: \**P* < 0.05, \*\**P* < 0.01, \*\*\**P* < 0.001, \*\*\*\**P* < 0.0001.

### Intratumoral injection of IFNγ-FasICD mRNA-LNP mediates regression of tumors in mice

To evaluate the effect of IFNγ–FasICD on tumor growth, syngeneic mice were subcutaneously inoculated with B16OVA melanoma cells. When tumors reached a size of 30–60 mm^3^, mice received intratumoral injections of either IFNγ–FasICD mRNA encapsulated in lipid nanoparticles (LNPs) or a control LNP formulation containing luciferase (Luc) mRNA (Figure 3A). Injections were administered five times at 2-day intervals, and tumor growth was monitored for approximately 20 days by measuring tumor volume. Tumors treated with IFNγ–FasICD mRNA-LNP exhibited a significant delay in tumor growth compared with those treated with Luc mRNA-LNP (n = 10) (Figures 3A, B). Consistent with these findings, all mice in the Luc mRNA-LNP control group died within 30 days, whereas 40% of mice treated with IFNγ–FasICD mRNA-LNP survived beyond this time point (Figure 3C). Comparable results were obtained in a second syngeneic tumor model using MC38 colon carcinoma cells. Intratumoral administration of IFNγ–FasICD mRNA-LNP markedly suppressed tumor growth relative to control treatment (Figures 3D, E). While all mice receiving Luc mRNA-LNP died within about 30 days, 20% of mice treated with IFNγ–FasICD mRNA-LNP remained alive over the same period (Figure 3F). Together, these results demonstrate that intratumoral delivery of IFNγ–FasICD mRNA-LNP effectively suppresses tumor progression and promotes tumor regression in multiple murine cancer models.

**Figure 3.**
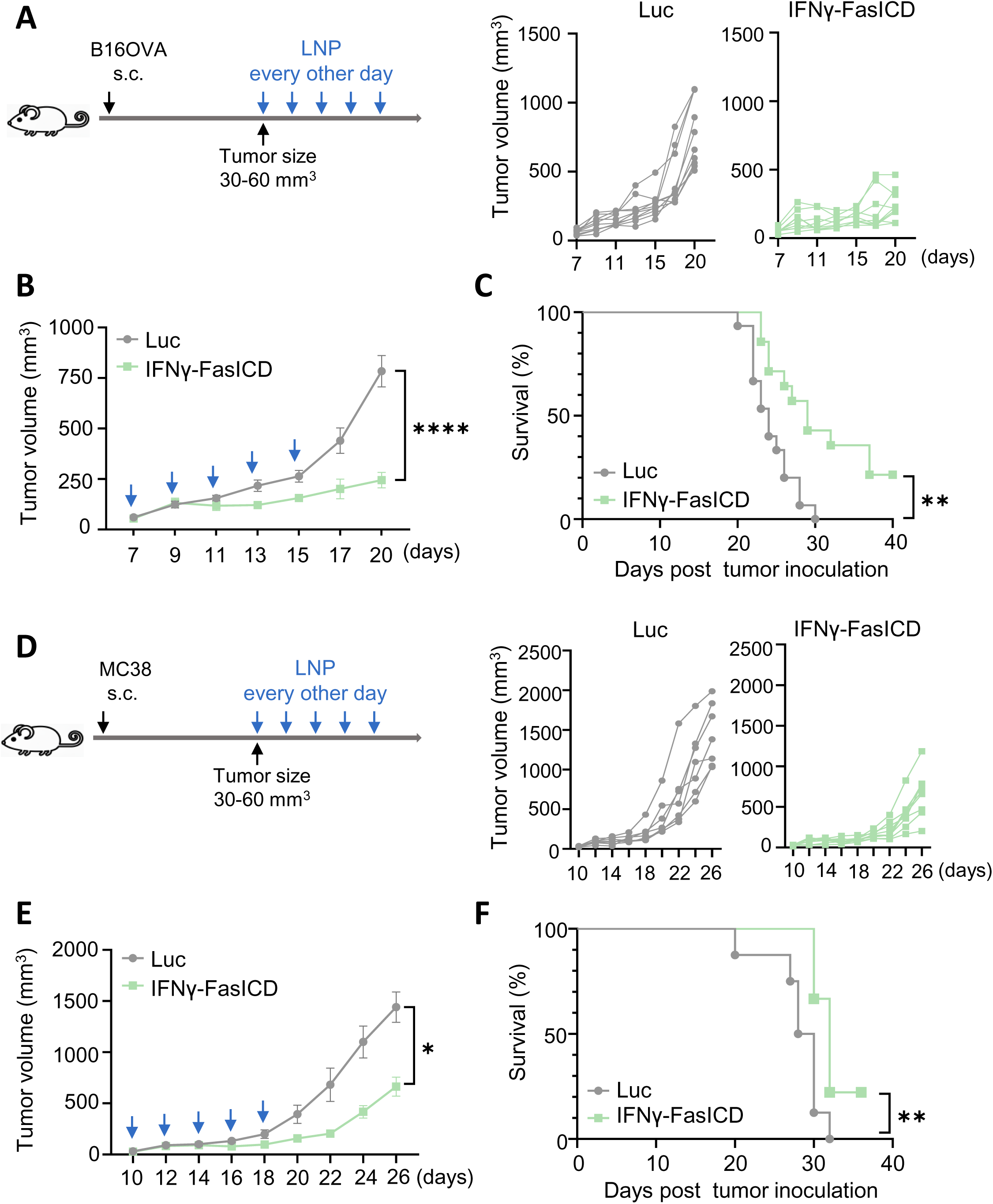
Intratumoral administration of IFNγ–FasICD LNP induces tumor regression. (**A**) Treatment schedule for the B16OVA tumor model in C57BL/6J mice. Mice were intratumorally injected with LNPs containing IFNγ-FasICD or luciferase control mRNA (10 µg mRNA /mouse) five times at 2-day intervals (left). Individual tumor growth curves for mice treated with luciferase (Luc) control or IFNγ-FasICD LNP are shown (*n* = 10) (right). **(B)** Mean tumor growth curves of B16OVA tumors. **(C)** Survival curves of mice bearing B16OVA tumors. **(D)** Treatment schedule for the MC38 tumor model in C57BL/6J mice (left). Individual tumor growth curves for mice treated with luciferase (Luc) control or IFNγ-FasICD LNP are shown (*n* = 7) (right). **(E)** Mean tumor growth curves of MC38 tumors. **(F)** Survival curves of mice bearing MC38 tumors. Statistical significance was determined by two-way ANOVA followed by Šídák’s multiple comparisons test **(B, E)**, and by the Mantel–Cox (log-rank) test **(C, F)**. Error bars indicate SEM. Note: \**P* < 0.05, \*\**P* < 0.01, \*\*\**P* < 0.001, \*\*\*\**P* < 0.0001.

### Intratumoral IFNγ–FasICD mRNA-LNP treatment promotes immune cell infiltration and functional activation of innate and adaptive immune cells in the tumor microenvironment

Previous studies have shown that apoptotic tumor cell death within the TME enhances dendritic cell infiltration and CD86 upregulation, thereby supporting CD8^+^ T cell priming and reinforcing local effector activity (51, 52). Therefore, we hypothesized that IFNγ–FasICD could similarly shape the TME by promoting immune cell recruitment and activation, thereby enhancing anti-tumoral immunity. As shown in Figure 4A, intratumoral administration of IFNγ–FasICD mRNA-LNP markedly increased the number of CD45^+^ immune cells by approximately 1.5-fold. In addition, CD8^+^ T cells were increased by 2-fold, while NK cells, NKT cells, conventional dendritic cells type 1 (cDC1s), and cDC2s were increased by more than 2-fold. Both M1 and M2 macrophages were also increased in the TME of IFNγ–FasICD–injected B16OVA tumors; however, the increase in M2 macrophages did not reach statistical significance (Figure 4A). In line with these increases in immune cell populations, IFNγ–FasICD administration significantly increased CD86 expression on both cDC1s and cDC2s, as well as NKG2D expression on NK and NKT cells within the TME, indicating their activation (Figure 4B). In parallel with cDC1 activation, the frequency of OVA-specific CD8^+^ T cells increased approximately 1.8-fold, although this increase did not reach statistical significance (Figure 4C). Notably, the frequency of FOXP3^+^ regulatory T cells was significantly reduced following IFNγ–FasICD injection. Consequently, the ratio of FOXP3^+^ T reg cells to OVA-specific CD8^+^ T cells was reduced by approximately 2.5-fold compared with the control group. To assess the maximal functional potential of CD8^+^ T cells and NK cells in the TME, *ex vivo* stimulation was performed using phorbol 12-myristate 13-acetate (PMA) and the calcium ionophore ionomycin. As shown in Figure 4D, IFNγ–producing T cells and NK cells were increased by approximately 2-fold and 2.5-fold, respectively, in IFNγ–FasICD–treated tumors compared with controls following *ex vivo* stimulation. Furthermore, the frequency of activated and proliferating CD8^+^ T cells (CD25^+^Ki-67^+^) was significantly increased. Taken together, these results indicate that IFNγ–FasICD remodels the TME toward a more immunostimulatory and anti-tumoral state, consistent with the tumor regression observed following intratumoral IFNγ–FasICD administration (Figure 3).

**Figure 4.**
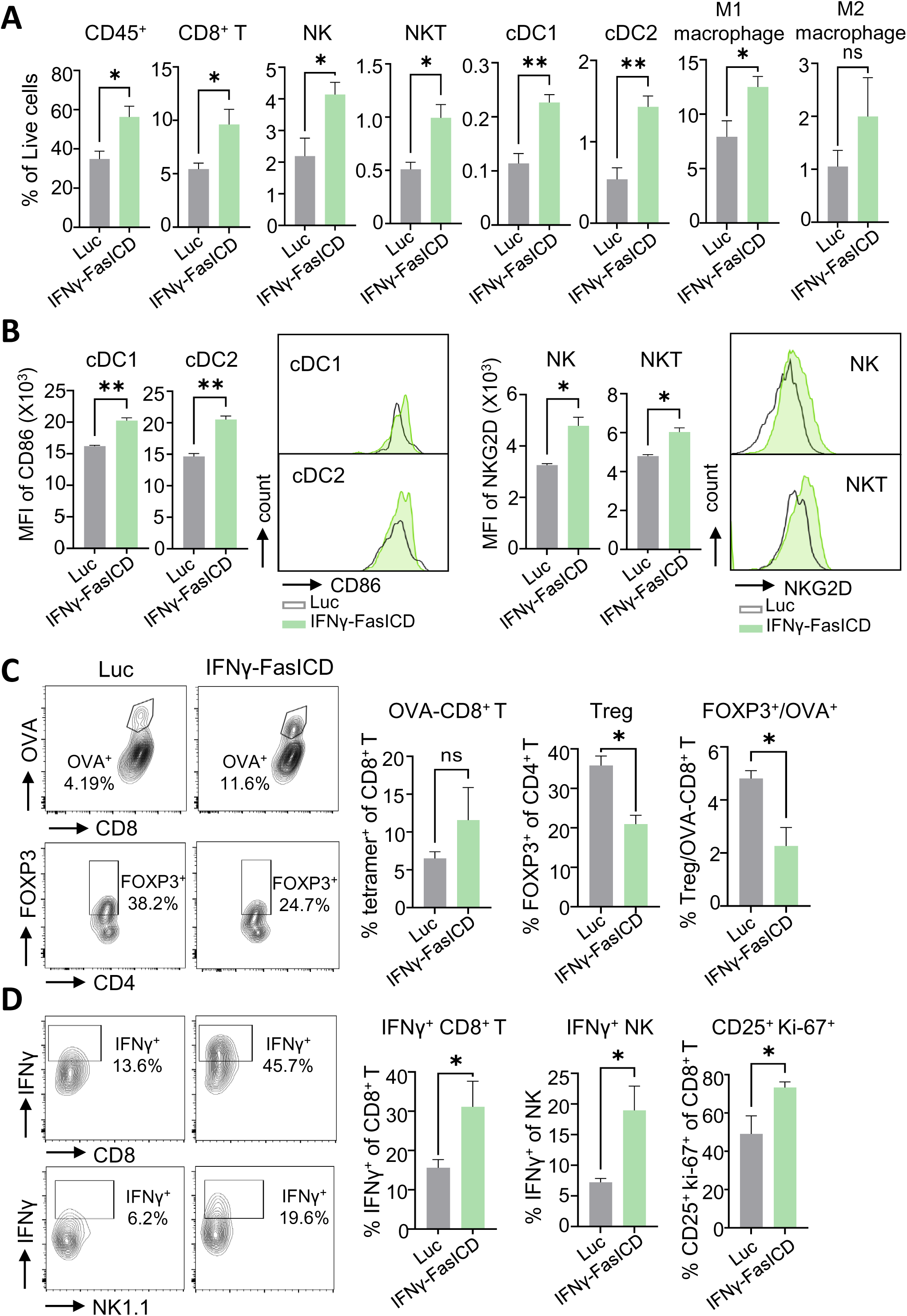
IFNγ-FasICD LNP treatment induces antitumoral immune responses in the B16OVA tumor microenvironment. Tumor-infiltrating immune cells within B16OVA tumors were analyzed by flow cytometry 3-9 days after the final intratumoral injection of control or IFNγ-FasICD LNP (*n* = 5), using the gating strategy shown in Supplementary Figure 2. **(A)** Frequencies of live CD45^+^ cells, CD8^+^ T cells, NK cells, NKT cells, conventional dendritic cell type 1 (cDC1s), cDC2, M1 macrophages, and M2 macrophages. **(B)** CD86 expression in cDC1 and cDC2 and NKG2D expression in NK and NKT cells are shown as bar graphs and representative histograms (MFI). **(C)** Representative flow cytometry plots show OVA-specific CD8^+^ T cells and FOXP3^+^ regulatory T cells (Tregs) (left). Mean frequencies and the ratio of FOXP3^+^ Tregs to OVA-specific CD8^+^ T cells are shown (right). **(D)** For intracellular cytokine analysis, cells were stimulated *ex vivo* with PMA and ionomycin for 6 h in the presence of Golgi inhibitors prior to staining. Representative plots and quantification of IFNγ^+^ CD8^+^ T cells, IFNγ^+^ NK cells, and CD25^+^Ki-67^+^ CD8^+^ T cells are shown. Statistical significance was determined by unpaired t-test. Error bars indicate SEM. Note: \**P* < 0.05, \*\**P* < 0.01, \*\*\**P* < 0.001, \*\*\*\**P* < 0.0001.

### IFNγ–FasICD increases dendritic cell migration and enhances effector potential of CD8^+^ T cells and NK cells across multiple immune compartments

Given the expansion of CD8^+^ T cells observed in IFNγ–FasICD-treated tumors, we next examined whether IFNγ–FasICD administration affected dendritic cell migration to tumor-draining lymph nodes (TdLNs). We found that intratumoral administration of IFNγ–FasICD mRNA-LNP increased the number of migratory dendritic cells by approximately 2.2-fold in TdLNs (Figure 5A). This increase was accompanied by a decrease in naïve CD8^+^ T cells (CD44^-^CD62L^+^) and a corresponding increase in the proportion of double-negative (DN; CD44^-^CD62L^-^) and central memory (CD44^+^CD62L^+^) CD8^+^ T cell subsets relative to the naïve population (Figure 5A). In peripheral compartments, increased frequencies of CD8^+^ T cells were observed in peripheral blood mononuclear cells (PBMCs), while significantly increased frequencies of NK cells were detected in both PBMCs and the spleen (Figures 5B, C). These immune changes were further associated with increased frequencies of IFNγ-producing CD8^+^ T cells in tumor-draining lymph nodes, the spleen, and PBMCs, as well as elevated frequencies of cytotoxic (GzmB^+^CD107a^+^) NK cells in PBMCs (Figures 5D, E). Together, these data indicate that IFNγ–FasICD treatment is associated with enhanced dendritic cell maturation and migration, accompanied by increased functional competence of CD8^+^ T cells and NK cells.

**Figure 5.**
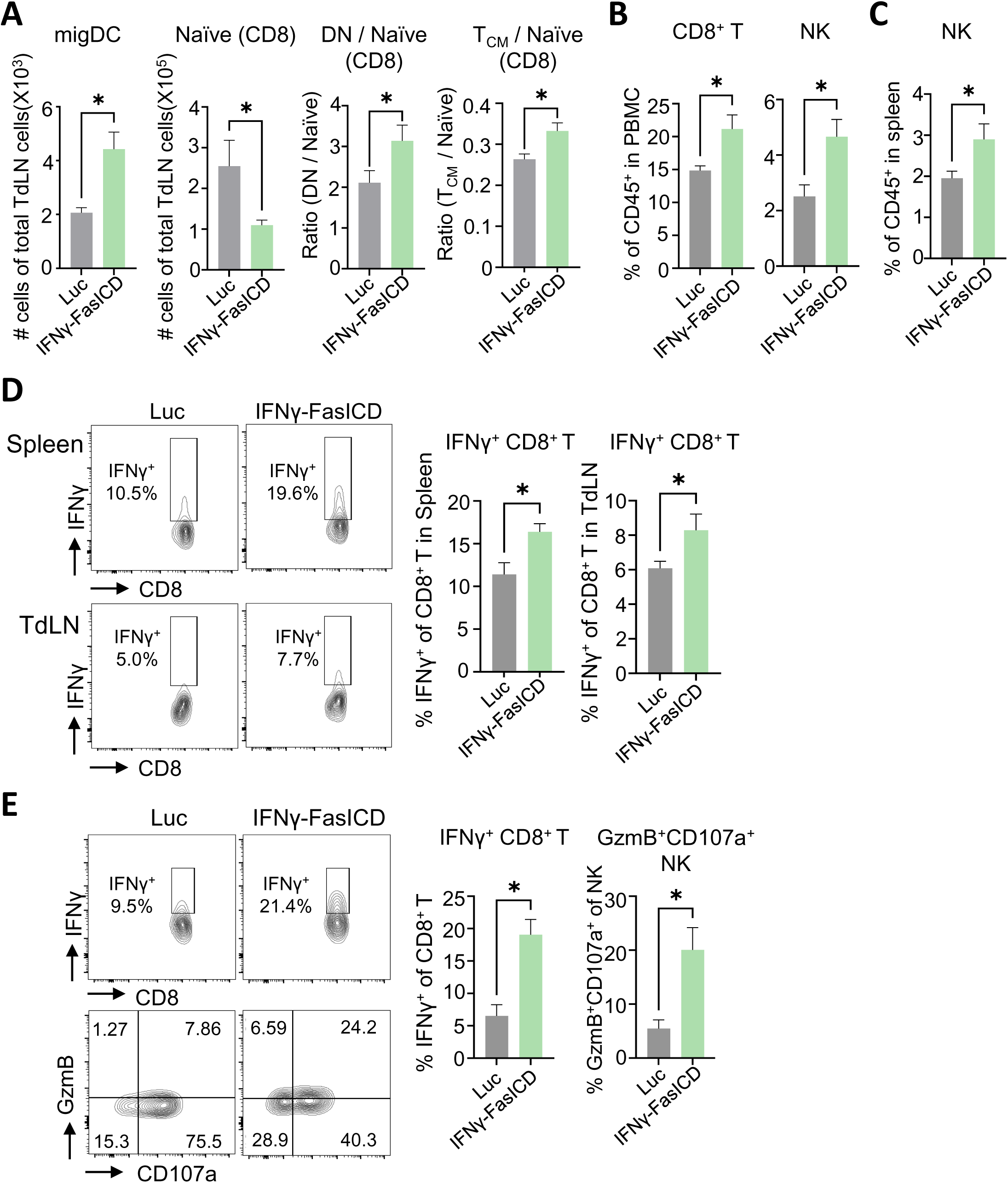
IFNγ–FasICD LNP promotes dendritic cell migration and enhances systemic effector activity of CD8^+^ T cells and NK cells. Immune cells in tumor-draining lymph nodes (TdLN), spleen, and peripheral blood mononuclear cells (PBMC) were analyzed by flow cytometry 3-9 days after the final intratumoral injection of control or IFNγ-FasICD LNP (*n* = 5), using the gating strategies shown in Supplementary Figures 3 and 4. **(A)** Numbers of migratory dendritic cells (migDCs), naïve CD8^+^ T cells, and the ratios of central memory CD8^+^ T cells (T^CM^) or double-negative (DN) CD8^+^ T cells to naïve CD8^+^ T cells in TdLN. Naive, CD44^-^CD62L^+^; T^CM^, CD44^+^CD62L^+^; DN, CD44^-^CD62L^-^. Absolute cell numbers were determined using an automated cell counter. **(B)** Frequencies of NK cells in the spleen. **(C)** Frequencies of CD8^+^ T cells and NK cells in PBMC. **(D, E)** For intracellular cytokine analysis, cells were stimulated *ex vivo* with PMA and ionomycin for 6 h in the presence of Golgi inhibitors prior to staining. **(D)** Representative flow cytometry plots and quantification of IFNγ^+^ CD8^+^ T cells in the spleen and TdLN. **(E)** Representative flow cytometry plots and quantification of IFNγ^+^ CD8^+^ T cells and Granzyme B^+^CD107a^+^ (GzmB^+^CD107a^+^) NK cells in PBMC. Statistical significance was determined by unpaired t-test. Error bars indicate SEM. Note: \**P* < 0.05, \*\**P* < 0.01, \*\*\**P* < 0.001, \*\*\*\**P* < 0.0001.

## Discussion

Cancer immunotherapy has been developed to target various mechanisms of immune cell activation. However, despite effective activation of cytotoxic immune cells, tumors frequently recur or metastasize, in part due to insufficient tumor antigen availability to sustain durable antitumor immune responses (53). In this study, we designed a membrane-anchored IFNγ–FasICD fusion protein to simultaneously engage interferon signaling and death receptor–mediated apoptosis in tumor cells. Intratumoral delivery of IFNγ–FasICD induces Fas-mediated tumor cell death and antigen release, thereby enhancing IFNγ–driven activation of local T cells and NK cells.

As shown in Figures 1 and 2, transfection of cancer cells with IFNγ–FasICD mRNA resulted in robust expression of IFNγ on the cell surface, accompanied by partial proteolytic release of soluble IFNγ, thereby enabling both membrane-restricted and diffusible cytokine activity. Functionally, IFNγ–FasICD expression induced marked tumor cell death *in vitro*, which was predominantly apoptotic rather than necrotic, as evidenced by caspase-3/7 activation and Annexin V staining. Comparison with the IFNγ–FasΔ construct demonstrated that IFNγ alone could induce partial cytotoxicity, consistent with its known context-dependent pro-death effects; however, inclusion of the Fas intracellular domain substantially amplified caspase-dependent apoptosis, indicating that FasICD provides a dominant and mechanistically distinct death signal. In parallel, IFNγ derived from IFNγ–FasICD–expressing tumor cells activated canonical IFNγ receptor signaling in both tumor cells and immune cells, as shown by induction of *Irf1* and *Cd274* in IFNγR-expressing tumor cells and STAT1 phosphorylation in ex vivo co-cultured splenocytes. Together, these data establish that the IFNγ–FasICD fusion protein operates through dual, non-redundant mechanisms: direct induction of apoptotic tumor cell death via Fas signaling and autocrine/paracrine activation of IFNγ pathways that are central to antitumor immune modulation. Therefore, this dual-function design has important conceptual implications for cancer gene therapy.

Our previous study demonstrated that CT26 stable cell lines expressing a membrane-bound IFNγ-Fas fusion protein, compared with IFNγ alone, functioned as an effective cancer vaccine, inducing systemic antitumor immunity mediated by T cells and NK cells (48). Building on these findings, we generated IFNγ–FasICD mRNA encapsulated in lipid nanoparticles (LNPs) and evaluated its therapeutic efficacy in B16OVA and MC38 tumor models. In vivo administration of IFNγ–FasICD mRNA-LNP, compared with the Luc mRNA-LNP control, markedly suppressed tumor growth and prolonged survival across two syngeneic tumor models, supporting its robust antitumor efficacy (Figure 3). IFNγ–FasICD treatment was associated with increased immune cell infiltration and a coordinated expansion of antigen-presenting cell populations together with cytotoxic lymphocytes (Figure 4). Notably, this immune remodeling included an expansion of OVA-specific CD8^+^ T cells, together with an increased frequency of activated and proliferating CD8^+^ T cells, as indicated by CD25 and Ki-67 expression, alongside enhanced conventional dendritic cell maturation and NK/NKT cell activation. Collectively, these results are consistent with strengthened interactions between APCs and T cells and the development of a pro-inflammatory immune microenvironment with antitumoral features. Consistent with these intratumoral changes, local immune remodeling was accompanied by enhanced migration of mature dendritic cells to tumor-draining lymph nodes, together with coordinated differentiation of CD8^+^ T cells toward pre-effector-like and memory phenotypes, including a double-negative (CD44^-^CD62L^-^) CD8^+^ T cell population previously defined as an intermediate pre-effector–like state arising during differentiation from naïve CD8^+^ T cells toward effector/memory fates (54) (Figure 5). In parallel, increased frequencies of IFNγ-producing CD8^+^ T cells and cytotoxic NK cell populations with increased Granzyme B and CD107a expression were observed in peripheral compartments. These results suggest a role for IFNγ–FasICD signaling in extending immune activation beyond the tumor microenvironment to regional and peripheral immune compartments.

Despite the robust antitumor efficacy observed *in vivo*, the activity of IFNγ–FasICD varies across tumor models, reflecting differences in intrinsic sensitivity to IFNγ–Fas signaling. *In vitro*, this variability is exemplified by the differential responses observed between B16OVA and MC38 tumor cells. Such variability may reflect intrinsic differences in IFNγ receptor expression and key components of the Fas signaling pathway, including FADD and c-FLIP (55–57). Moreover, beyond direct effects on tumor cells, IFNγ can differentially influence NK cell–mediated cytotoxicity by modulating the expression of activating and inhibitory NK cell ligands in a tumor type–dependent manner (58). Taken together, these observations indicate that the responsiveness to IFNγ–FasICD is shaped by intrinsic molecular features of specific tumor types and should be taken into account in the assessment of Fas- and IFNγ-based therapeutic strategies.

The immunological effects of IFNγ–FasICD *in vivo* extend beyond CD8^+^ T cell responses and arise from the coordinated action of IFNγ signaling and Fas-mediated tumor cell death across multiple immune compartments, rather than from IFNγ signaling alone. IFNγ can contribute to the maturation and migratory programming of dendritic cells, but these effects emerge most robustly in contexts where additional signals, such as TLR-mediated innate immune stimulation, are present (59, 60). Moreover, IFNγ can indirectly support the upregulation of NKG2D through an IL-12–dependent feedback loop, in which intratumoral conventional dendritic cells produce IL-12 in response to IFNγ while concurrently undergoing maturation via increased expression of costimulatory molecules (61–63). In this context, Fas-mediated apoptotic tumor cells generated in the presence of IFNγ may provide a complementary maturation signal to dendritic cells, thereby enhancing CD86 expression and promoting the accumulation of migratory dendritic cells in tumor-draining lymph nodes. These functionally activated dendritic cells may subsequently facilitate NKG2D upregulation on NK and NKT cells, consistent with prior reports indicating that IFNγ alone is insufficient to fully induce this activation pathway (64). IFNγ–FasICD treatment was also associated with marked alterations in CD8^+^ T cell subset composition within tumor-draining lymph nodes, including a reduction in naïve T cells and a concomitant increase in pre-effector-like and central memory populations. These changes are consistent with enhanced T cell priming and differentiation, driven by increased antigen availability following Fas-mediated tumor cell death, together with the accumulation of migratory dendritic cells capable of efficient antigen cross-presentation in tumor-draining lymph nodes.

IFNγ–FasICD may also overcome key limitations associated with therapeutic strategies that rely solely on full-length Fas overexpression. Fas signaling is often ineffective within the tumor microenvironment due to the limited availability of Fas ligand (FasL). Recent studies indicate that FasL expression is largely restricted to T cells and NK cells and is minimal in tumor or stromal cells from cancer patients. Moreover, reduced infiltration or functional impairment of these immune populations can further constrain the efficacy of Fas-based approaches (65). In our system, IFNγ–FasICD bypasses the requirement for endogenous FasL, by inducing Fas-mediated apoptotic signaling without dependence on FasL availability. Nevertheless, restoration of apoptotic signaling alone may not be sufficient for therapeutic benefit. Tumor cell apoptosis—whether constitutive or induced by radiotherapy or chemotherapy—has been reported to paradoxically promote protumoral polarization of tumor-associated macrophage (TAMs) (66–68). In our study, although M2 macrophages were modestly increased, this change did not reach statistical significance, whereas M1 macrophages were significantly expanded, resulting in a tumor microenvironment skewed toward an antitumoral phenotype (Figure 4A). These findings suggest that IFNγ–FasICD not only induces tumor cell apoptosis but also avoids the adverse macrophage reprogramming often associated with apoptotic tumor debris. Consequently, IFNγ–FasICD may offer a distinct advantage over Fas-based monotherapy by preserving or enhancing M1 macrophage polarization while simultaneously promoting NK cell activation and CD8^+^ T cell effector function within the tumor microenvironment. Additionally, IFNγ has been reported to antagonize immunosuppressive pathways, including the inhibition of regulatory T cell (Treg) differentiation (69). Although IFNγ-dependent enhancement of Treg activity has been described in certain chronic inflammatory settings (70, 71), such effects were not dominant in our model, as evidenced by a significant reduction in FOXP3^+^ Treg populations. This observation is consistent with a net immunostimulatory outcome driven by IFNγ–FasICD treatment.

Consistent with our findings, several studies have reported enhanced antitumor efficacy when tumor cell death–inducing strategies are combined with immunotherapy. Approaches such as the combination of radiotherapy with immune checkpoint blockade, including anti-PD-L1 therapy, have been actively explored, with ionizing radiation shown to exert antitumor effects largely through CD8^+^ T cell–dependent mechanisms (72–74). Moreover, accumulating evidence indicates that tumor cell–intrinsic alterations in death receptor signaling can influence susceptibility to chimeric antigen receptor (CAR) T cell–mediated cytotoxicity (75). In this context, IFNγ–FasICD treatment could sensitize tumor cells to CAR T cell–mediated killing by enabling Fas ligand–independent engagement of downstream apoptotic machinery, including FADD–CASP8 axis signaling, or by lowering the apoptotic threshold through stabilization of downstream death receptor signaling components. Moreover, IFNγ signaling has the potential to upregulate additional death receptors, such as TNFR and TRAIL receptors, thereby expanding the repertoire of death receptor–mediated pathways available for CAR T cell–induced cytotoxicity and supporting potential combinatorial effects. In addition, Fas–FasL signaling has been implicated in both antigen-specific and localized bystander tumor killing, and IFNγ exposure has been shown to increase surface Fas expression, potentially further augmenting this process (76). Collectively, these observations suggest that tumor-targeted IFNγ–FasICD–based strategies may serve as a rational and mechanistically complementary adjunct to T cell–directed immunotherapies.

IFNγ is well recognized for its pleiotropic roles in tumor biology, exerting both tumor-suppressive and tumor-promoting effects depending on dose, duration, and cellular context (77–79). By tethering IFNγ to the tumor cell membrane while simultaneously coupling it to Fas-mediated apoptosis, our strategy spatially restricts IFNγ activity to the tumor microenvironment and links cytokine signaling to irreversible cell-intrinsic death pathways. This design may mitigate systemic toxicity associated with IFNγ while favoring immunogenic apoptosis over necrosis, a distinction that is increasingly appreciated as critical for productive antitumor immunity. Our findings, therefore, provide a mechanistic foundation for the *in vivo* antitumor effects observed and support the broader concept that rationally engineered cytokine–death receptor fusion proteins can reprogram tumor cells into localized sources of both apoptotic signals and immune activation cues.

## Material and methods

### Tumor Cell Lines and Mice

Female C57BL/6J mice, aged 6-8 weeks, were purchased from Hana Biotech (Korea). All procedures involving animals were approved by the Institutional Animal Care and Use Committee (IACUC) of Pohang University of Science and Technology (POSTECH-2025-0038) and conducted in accordance with institutional guidelines. HEK293T cells were obtained from ATCC (CRL-3216) HEK293T, MC38, B16OVA cells were cultured in DMEM (Welgene, LM001-05, Korea) supplemented with 10% heat-inactivated fetal bovine serum (FBS; Atlas Biologicals) and 1% penicillin-streptomycin (10,000 U/mL, Gibco) in a humidified atmosphere with 5% CO^2^ at 37°C.

### Antibodies

Brilliant Violet 605 anti-mouse CD45 Antibody (BioLegend, 103139), Brilliant Violet 650 anti-mouse/rat XCR1 Antibody (BioLegend, 148220), BUV395 Rat Anti-Mouse CD172a (BD, 740282), Brilliant Violet 785 anti-mouse NK-1.1 Antibody (BioLegend, 108749), FITC Hamster Anti-Mouse CD11c (BD, 553801), PE Hamster Anti-Mouse CD11c (BD, 553802), PerCP/Cyanine5.5 anti-mouse I-A/I-E Antibody (BioLegend, 107625), PE anti-mouse CD314 (NKG2D) Antibody (BioLegend, 130207), PE/Cyanine7 anti-mouse CD86 Antibody (BioLegend, 105013), Alexa Fluor 700 anti-mouse CD317 (PDCA-1) Antibody (BioLegend, 127038), APC/Cyanine7 anti-mouse CD3 Antibody (BioLegend, 100222), FITC Rat Anti-Mouse CD4 (BD, 553729), PE/Cyanine7 anti-mouse CD8a Antibody (BioLegend, 100721), FITC anti-mouse CD206 (MMR) Antibody (BioLegend, 141703), APC anti-mouse F4/80 Antibody (BioLegend, 123116), PerCP/Cyanine5.5 anti-mouse/human CD11b Antibody (BioLegend, 101228), FITC anti-mouse I-A/I-E Antibody (BioLegend, 107605), PerCP/Cyanine5.5 anti-mouse CD103 Antibody (BioLegend, 121415), PE Rat Anti-Mouse CD25 (BD, 553866), APC anti-mouse CD8a Antibody (BioLegend, 100712), PE Rat Anti-Mouse CD44 (BD, 553134), APC anti-mouse CD62L Antibody (BioLegend, 104411), PE/Cyanine7 anti-mouse CD107a (LAMP-1) Antibody (BioLegend, 121619), PE anti-mouse CD274 (PD-L1) Antibody (BioLegend, 124307), PE IFN-γ (BD, 554412), PerCP/Cyanine5.5 Granzyme B Antibody (BioLegend, 372211), APC FOXP3 Antibody (Invitrogen, 17-5773-80), PE/Cyanine7 Ki-67 Antibody (BioLegend, 151217) and PE Arginase 1(ARG1) Antibody (R&D systems, IC5868P) were used.

### RNA Extraction and qPCR

For detection of *Ifngr1* and *Ifngr2* expression in B16OVA and MC38 cell lines, total RNA was extracted using TRIzol (GeneAll, 301-001, Korea). RNA concentrations were measured using an Ultrospec 8000 spectrophotometer (GE Healthcare). Total RNA (2 µg) was reverse transcribed using the AccuPower RT Premix (Bioneer, K-2041, Korea). Semi-quantitative reverse transcription polymerase chain reaction (RT-PCR) was performed using Taq polymerase (Genetbio, G-2000, Korea) and the appropriate primers (Supplementary Table 1). The mRNA levels of *Irf1* and *Cd274* in B16OVA and MC38 cells following *in vitro* mRNA transfection were analyzed by real-time quantitative RT-PCR (qRT-PCR) on the QuantStudio 3 Real-Time PCR System (Thermo Fisher Scientific) using a 2X Real-Time PCR Master Mix containing SFCgreen (BioFACT, DQ362-40h, Korea) and the appropriate primers (Supplementary Table 1). 18S rRNA was used for normalization in the qRT-PCR analysis. Each PCR reaction was performed in triplicate. Relative mRNA levels were calculated using the 2 ^(-ΔΔCt)^ method.

### Enzyme-Linked Immunosorbent Assay (ELISA)

The expression level of IFNγ in HEK293T cells was quantified using an IFNγ ELISA kit (Invitrogen, 88-7314). Cells were seeded in 24-well plates and were transfected the following day with 0.25 µg of mRNA in 100 µL Opti-MEM using Lipofectamine 2000. The total volume was adjusted to 400 µL per well with culture medium. Six hours after transfection, both culture supernatants and cell lysates were collected. Total cell lysates (50 µL) and supernatants (50 µL) were diluted according to the manufacturer’s instruction prior to ELISA analysis.

### MTT assay

Cells were plated in 48-well culture plates and were transfected the following day with 0.25 µg of mRNA using Lipofectamine 2000. After 24 hours, cell viability was assessed using the MTT assay (3-(4,5-dimethylthiazol-2-yl)-2,5-diphenyltetrazolium bromide; GoldBio, T-030-1, USA).

### Caspase-3/7 activity

Caspase-3/7 activity was assessed using a kit purchased from Promega (G8090) according to the manufacturer’s instructions. Cells were seeded in 96-well culture plates and were transfected the following day with 50 ng of mRNA using Lipofectamine 2000. Four hours after transfection, the medium was replaced with fresh culture medium (50 µL per well), and cells were incubated at 37°C for 3 hours. Caspase-3/7 reagent was added, and the plates were incubated at room temperature for 90 minutes. The supernatants were collected and transferred to white-walled plates, and luminescence was measured using a plate-reading luminometer (Synergy HTX Multimode Reader, BioTek, USA).

### Analysis of Apoptosis by Annexin V/PI Staining

Cells were plated in 6-well culture plates and were transfected the following day with 1 µg of mRNA using Lipofectamine 2000. After 24 hours, apoptosis was assessed by staining APC-Annexin V (BD, 550474) and 10 µg/mL propidium iodide (PI; Sigma-Aldrich, P4170). Stained cells were analyzed using flow cytometry (CytoFLEX LX, Beckman Coulter), and data were processed using FlowJo software (BD).

### Tumor Challenge

For tumor implantation, 3 × 10^5^ to 5 × 10^5^ MC38 or B16OVA cells were injected subcutaneously into the right flank of female C57BL/6 mice in 100 µL of Dulbecco’s phosphate-buffered saline (DPBS; Welgene, LB 001-02). Tumor growth, body weight, and survival were monitored every other day. Tumor size was measured using calipers, and tumor volume was calculated using the formula 0.52 x S^2^ x L, where L is tumor length and S is tumor width. When tumor volumes reached approximately 30–60 mm³, mice were randomized into treatment groups to ensure comparable average tumor sizes and intratumoral injections of mRNA-LNPs were then administered. In accordance with the IACUC guidelines of Pohang University of Science and Technology, the humane endpoint was defined as a tumor volume exceeding 1,500 mm^3^.

### Tissue Dissociation

To analyze immune cells in tumor tissues, tumors were mechanically dissociated using a 3 mL syringe plunger and filtered through 70-μm cell strainers (SPL, 93070, Korea) in 1 X PBS. Following centrifugation, cells were incubated in a digestion buffer consisting of RPMI-1640 supplemented with 80 U/mL collagenase IV [Gibco, 17104-019] and 80 U/mL DNase I [Sigma-Aldrich, D5025] at 37°C for 20 minutes. The cells were then filtered through a 40-μm cell strainer (SPL, 93040) and enriched by OptiPrep (Sigma-Aldrich, D1556) density gradient separation according to the manufacturer’s instructions.

To assess immune cells in tumor-draining lymph nodes (TdLNs), TdLNs were collected and mechanically dissociated by passing them through cell strainers. For TdLN samples, total viable cell numbers were determined using an automated cell counter (LUNA-FL Automated Fluorescence Cell Counter, Logos Biosystems, Korea) according to the manufacturer’s instructions. For *in vitro* direct co-culture experiments, spleens from naïve mice were harvested, and single-cell suspensions were prepared by passing them through cell strainers. Red blood cells were removed from splenocyte preparations by incubation with RBC lysis buffer at 4°C for 10 minutes.

### Flow Cytometry Analysis

For surface staining, cells were blocked for Fcγ receptors (anti-mouse CD16/32; BioLegend, 156604) and stained with a viability dye (Violet 510 Ghost Dye; Tonbo, 13-0870-T100, USA), followed by staining with fluorophore-conjugated antibodies for 30 minutes at 4°C. For OVA-specific CD8^+^ T cell detection, cells were stained with a PE-conjugated H-2Kb/OVA (SIINFEKL) tetramer (ACROBiosystems, H2A-MP2H7, USA) according to the manufacturer’s protocol. For intracellular cytokine staining, cells were fixed and permeabilized using the Fixation/Permeabilization Kit (BD, 554714) according to the manufacturer’s instructions, and stained with antibodies against IFNγ and Granzyme B. For intracellular transcription factor staining, the Foxp3 Staining Buffer Kit (eBioscience, 00-5523-00) was used according to the manufacturer’s instructions, followed by staining with antibodies against FOXP3, Ki-67, or Arginase-1 (ARG1). For intracellular pSTAT1 staining, cells were fixed with 4% formaldehyde for 10 minutes at room temperature and washed. Cells were first stained with an anti-CD45 antibody to identify immune cells and were subsequently permeabilized with ice-cold methanol for 10 minutes at 4°C with gentle vortexing. After washing, cells were stained with an anti-pSTAT1 antibody (Cell Signaling, 9167) diluted in 1% BSA/PBS for 40 minutes at room temperature, followed by staining with an Alexa Fluor 488–conjugated anti-rabbit secondary antibody (Invitrogen, A-11034). Data were acquired on a CytoFLEX LX flow cytometer (Beckman Coulter).

### Confirmation of Membrane-Bound IFNγ Expression

To evaluate membrane-bound IFNγ expression on HEK293T cells, cells were plated in 6-well culture plates and were transfected the following day with 1 µg of mRNA using Lipofectamine 2000. After 24 hours, cells were detached using Accutase (Merck Millipore, SCR005) and stained for surface IFNγ. To assess membrane-bound IFNγ expression in tumor tissue, mRNA-LNPs were injected intratumorally when tumors reached approximately 80 mm³. At each designated time point, tumors were processed into single-cell suspensions using the previously described protocol, except for the OptiPrep step. Cells were stained with a viability dye, followed by surface staining with antibodies against CD45 and IFNγ as described above.

### Assessment of IFNγ-Induced STAT1 Phosphorylation in Splenocytes

B16OVA cells were plated in 24-well plates and transfected with 0.25 µg of mRNA. After 24 hours of transfection in complete medium (500 µL per well), 3 × 10^6^ naïve splenocytes were suspended in 500 µL medium without FBS and were directly co-cultured with the transfected B16OVA cells. Samples were then harvested, fixed and stained according to the previously described staining protocols.

### *Ex Vivo* Cytokine and Granzyme B Detection

Single-cell suspensions obtained from dissociated tumor tissue, tumor-draining lymph nodes, spleen, and PBMCs were cultured in complete RPMI-1640 medium in 96-well plates. Cells were stimulated with PMA/ionomycin (eBioscience, 00-4970-93) for 4 hours at 37°C in the presence of brefeldin A (Merck Millipore, B5936, 5 µg/mL), monensin (Sigma-Aldrich, M5273, 5 µg/mL), 50 µM β-mercaptoethanol, and an anti-CD107a antibody to monitor degranulation. After stimulation, cells were stained with a viability dye, followed by surface staining with antibodies against the appropriate markers. For intracellular cytokine staining, cells were fixed and permeabilized using the Fixation/Permeabilization Kit (BD, 554714) according to the manufacturer’s instructions, and stained with antibodies against IFNγ and granzyme B.

### mRNA-LNPs preparation

DNA constructs encoding codon-optimized IFNγ–FasICD, IFNγ–FasΔ, and firefly luciferase (Luc) were generated by cloning the respective open reading frames into a plasmid backbone derived from a SARS-CoV-2 mRNA vaccine (BNT162b2; Pfizer–BioNTech). The plasmid DNA was linearized at the end of the poly(A) tail using BsaI (New England Biolabs). The linearized plasmid DNA was used as template for *in vitro* transcription (IVT). The IVT reaction mixture included the linearized plasmid DNA, spermidine trihydrochloride (Sigma-Aldrich), magnesium chloride hexahydrate (Sigma-Aldrich), PEG8000 (Sigma-Aldrich), dithiothreitol (GoldBio), CleanCap AG (TriLink Biotechnologies, USA), ATP, GTP, and CTP (Thermo Fisher Scientific), N1-methyl-pseudouridine-5′-triphosphate (TriLink Biotechnologies), purified T7 RNA polymerase, RNase Inhibitor (Roche), Tris-HCl (pH 8.0) buffer in DEPC-treated distilled water. IVT reaction was performed at 37°C for 2 hours and followed by treatment with DNase I at 37°C for 20 minutes to degrade the template DNA. The IVT mRNA was purified using a Monarch Spin RNA Cleanup Kit (New England Biolabs). The purified mRNA was quantified using a NanoDrop One spectrophotometer (Thermo Fisher Scientific), and its integrity was assessed by agarose gel electrophoresis.

mRNA-LNPs were prepared by microfluidic mixing using a NanoAssemblr Spark or NanoAssemblr Ignite (Precision NanoSystems, Canada). mRNA dissolved in 50 mM citrate buffer (pH 4.0) was mixed with lipids at a volume ratio of 2:1 or 3:1 (mRNA:lipid) and an N/P ratio of 6 and at a total flow rate of 12 mL/min. The lipids, consisting of SM-102 (Echelon Biosciences, USA), DSPC (Sigma), cholesterol (Sigma), and DMG-PEG2000 (Avanti, USA) were dissolved in ethanol at a molar ratio of 50:10:38.5:1.5. Following formulation, mRNA-LNPs were buffer-exchanged into DPBS and concentrated using a 30 kDa MWCO Amicon Ultra centrifugal filter unit (Merck Millipore). The resulting mRNA-LNPs were mixed with sucrose to a final concentration of 8% (w/v) and stored at -20°C until use.

### mRNA-LNPs characterization

Particle size, polydispersity index (PDI) and zeta potential were measured using a Zetasizer Nano ZS (Malvern Panalytical, United Kingdom). Encapsulation efficiency was determined using a RediPlate™ 96 RiboGreen™ RNA Quantitation Kit (Thermo Fisher Scientific), according to the manufacturer’s protocol. Briefly, encapsulated mRNAs treated with or without 1% Triton X-100 were quantified (Q^triton treated^ and Q^triton untreated^, respectively) using a Synergy HTX (BioTek). The encapsulation efficiency (EE) was calculated as follows: EE (%) = (Q^triton treated^ - Q^triton untreated^)/Q^triton treated^ × 100. Encapsulation of mRNA-LNPs was further evaluated by 1.5% agarose gel electrophoresis. As a control for LNP disruption, a portion of the mRNA-LNPs was treated with Triton X-100 prior to loading. The RNA ladder and free mRNA controls were heated at 70°C for 10 minutes. After cooling, the mRNA-LNPs and heated free mRNA controls were mixed with loading dye, loaded onto the gel, and run in TAE buffer alongside the RNA ladder (Thermo Scientific). The morphology of mRNA-LNPs was examined by cryo-electron microscopy. Briefly, LNP samples were applied to glow-discharged Quanti-foil holey carbon grids (Quantifoil Micro Tools, Germany), blotted to remove excess liquid, and rapidly vitrified using a Vitrobot Mark IV (Thermo Fisher Scientific). Images were acquired using a Talos Glacios microscope (Thermo Fisher Scientific).

### Bioluminescence imaging

For *in vivo* bioluminescence imaging, Luc-encoding naked mRNA in PBS, naked mRNA in Ringer’s lactate, or Luc-encoding mRNA–LNPs were injected intratumorally when tumors reached approximately 320 mm³. At each designated time point, 200 µL of 15 mg/mL D-luciferin (Goldbio) in PBS was injected intraperitoneally. Bioluminescence was measured 10 minutes after injection of luciferin using the Ami HTX imaging system (Spectral Instruments Imaging, USA). Data analysis was performed using Aura software (Spectral Instruments Imaging).

### Statistical Analysis

All data are presented as mean ± SEM. Statistical analyses were performed using GraphPad Prism 9 (GraphPad Software, USA). Depending on the experimental design, t-tests, one-way ANOVA, or two-way ANOVA were applied to assess differences between groups (\**P* < 0.05, \*\**P* < 0.01, \*\*\**P* < 0.001, \*\*\*\**P* < 0.0001). Survival curves were generated using Kaplan–Meier method, and statistical differences between groups were assessed using the Mantel–Cox (log-rank) test in GraphPad Prism 9.

## Author Contributions

H.-S.S performed all experiments, analyzed the data, and wrote the paper. S.-G.K mainly focused on LNP-mRNA production, IV injection & analysis and wrote the paper. H.L.and J.-O.L. designed the research, analyzed the data, and wrote the paper. All authors have read and approved the final paper.

## ACKNOWLEDGMENTS

We thank Prof. Seung-Woo Lee of POSTECH, Pohang, for generously providing the MC38 and B16OVA cell lines used in this study.

