## Supplementary Figure 1-4 for "Integrating Fas-mediated apoptosis with IFNγ signaling to drive tumor regression in mRNA cancer therapeutics"

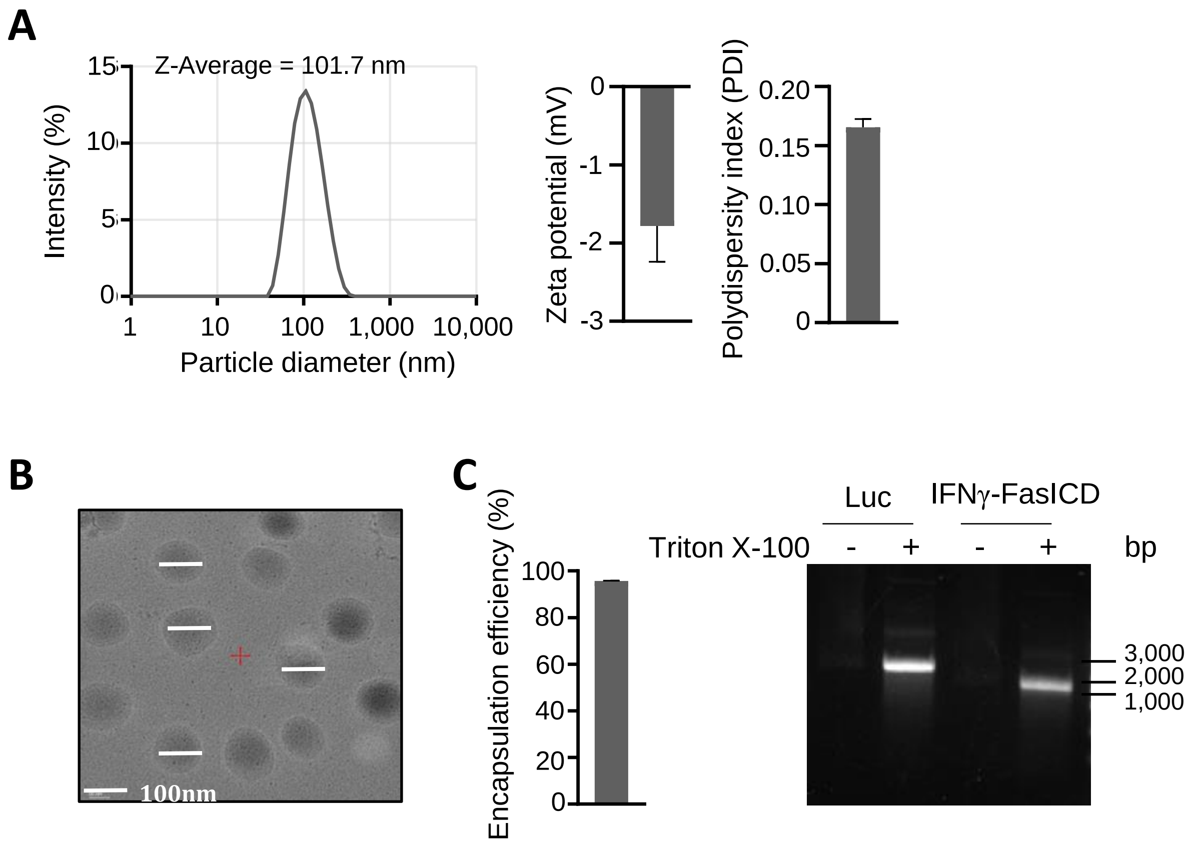


**Supplementary Figure 1. Characterization of LNP incorporating mRNA. (A)** size, zeta potential and polydispersity index (PDI) of LNP incorporating mRNA were measured using dynamic light scattering (DLS). **(B)** Images of LNP incorporating mRNA using cryo-electron microscopy. **(C)** Encapsulation efficiency was verified by Ribogreen assay (left) and gel electrophoresis with/without triton x-100 treatment (right).

### Supplementary Figure 2.

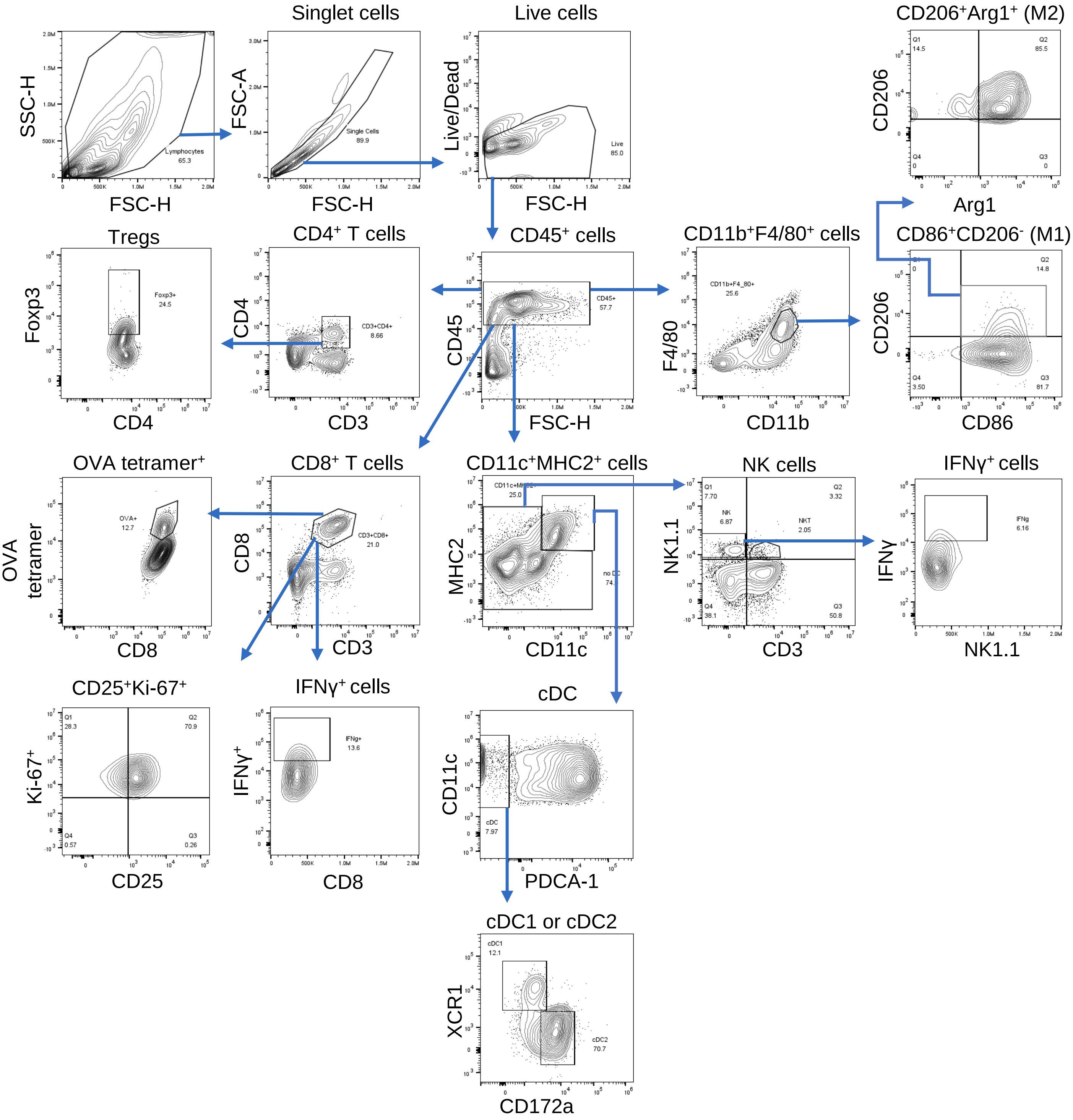


**Supplementary Figure 2. Flow cytometry gating strategy for the characterization of tumor-infiltrating immune cell subsets.** A detailed gating strategy for tumor-infiltrating lymphocytes (TILs), as shown in Figure 4. After live CD45$^{+}$ immune cells gating, each cell subset was gated as follows: M1 macrophage; CD11b$^{+}$F4/80$^{+}$CD86$^{+}$CD206$^{-}$, M2 macrophage; CD11b$^{+}$F4/80$^{+}$CD206$^{+}$Arg1$^{+}$, CD4$^{+}$ T; CD3$^{+}$CD4$^{+}$, Treg; CD3$^{+}$CD4$^{+}$Foxp3$^{+}$, CD8$^{+}$ T; CD3$^{+}$CD8$^{+}$, cDC1; CD11c$^{+}$MHC2$^{+}$PDCA-1$^{-}$XCR1$^{+}$CD172a$^{-}$, cDC2; CD11c$^{+}$MHC2$^{+}$PDCA-1$^{-}$ XCR1$^{-}$ CD172a$^{+}$, NK ; CD3$^{-}$ NK1.1$^{+}$

### Supplementary Figure 3.

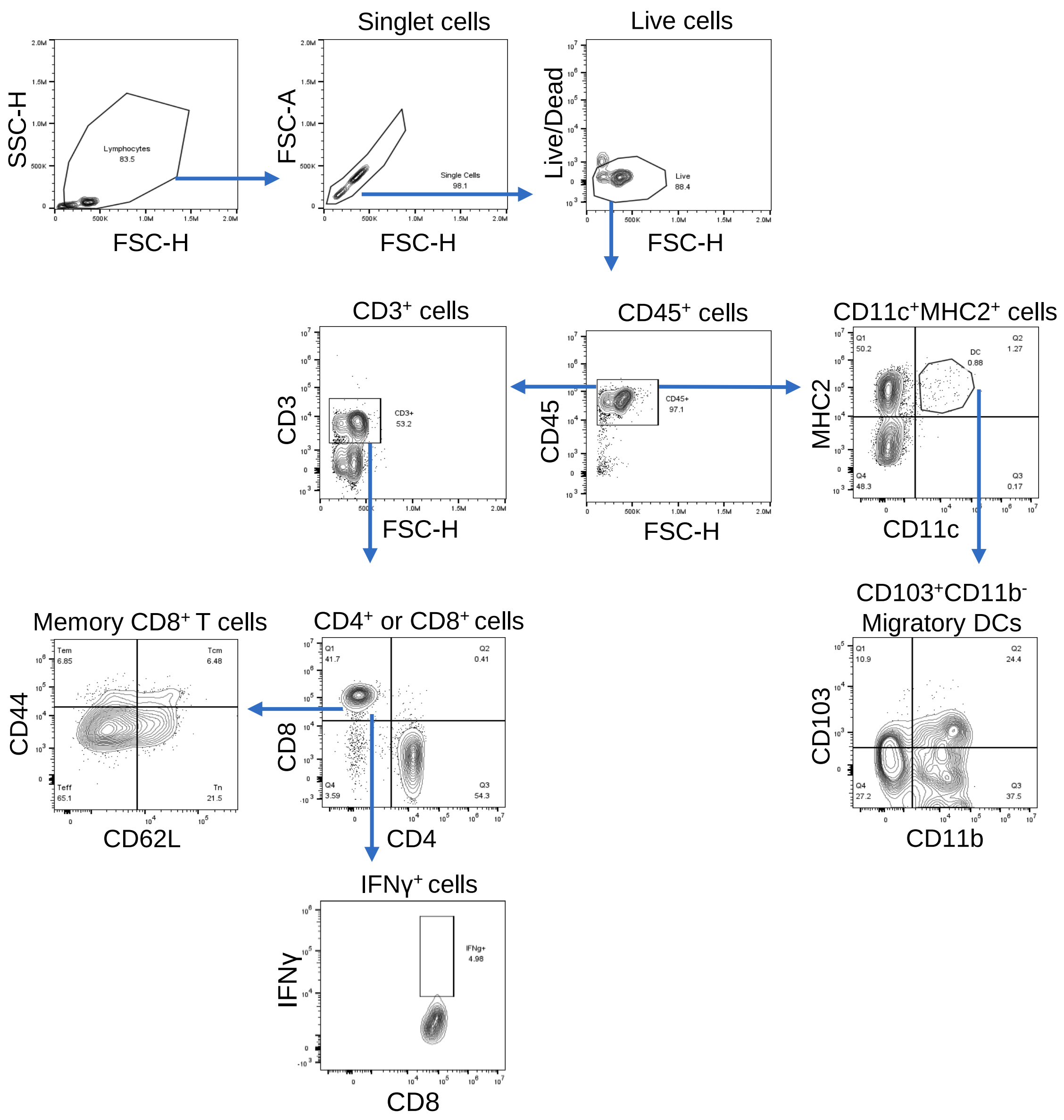


**Supplementary Figure 3. Flow cytometry gating strategy for the characterization of immune cell subsets in tumor-draining lymph nodes.** A detailed gating strategy for migratory DCs and memory T cells, as shown in Figure 5. After live CD45$^+$ immune cells gating, each cell subset was gated as follows: Naïve CD8$^+$ T cells; CD3$^+$CD8$^+$CD44$^-$CD62$^+$, DN CD8$^+$ T cells; CD3$^+$CD8$^+$CD44$^-$CD62$^-$, Central memory ($T_{CM}$) CD8$^+$ T cells; CD3$^+$CD8$^+$CD44$^+$CD62L$^+$, Effector memory ($T_{EM}$) CD8$^+$ T cells; CD3$^+$CD8$^+$CD44$^+$CD62$^-$, migratory DC; CD11c$^+$MHC2$^+$CD103$^+$CD11b$^-$

### Supplementary Figure 4.

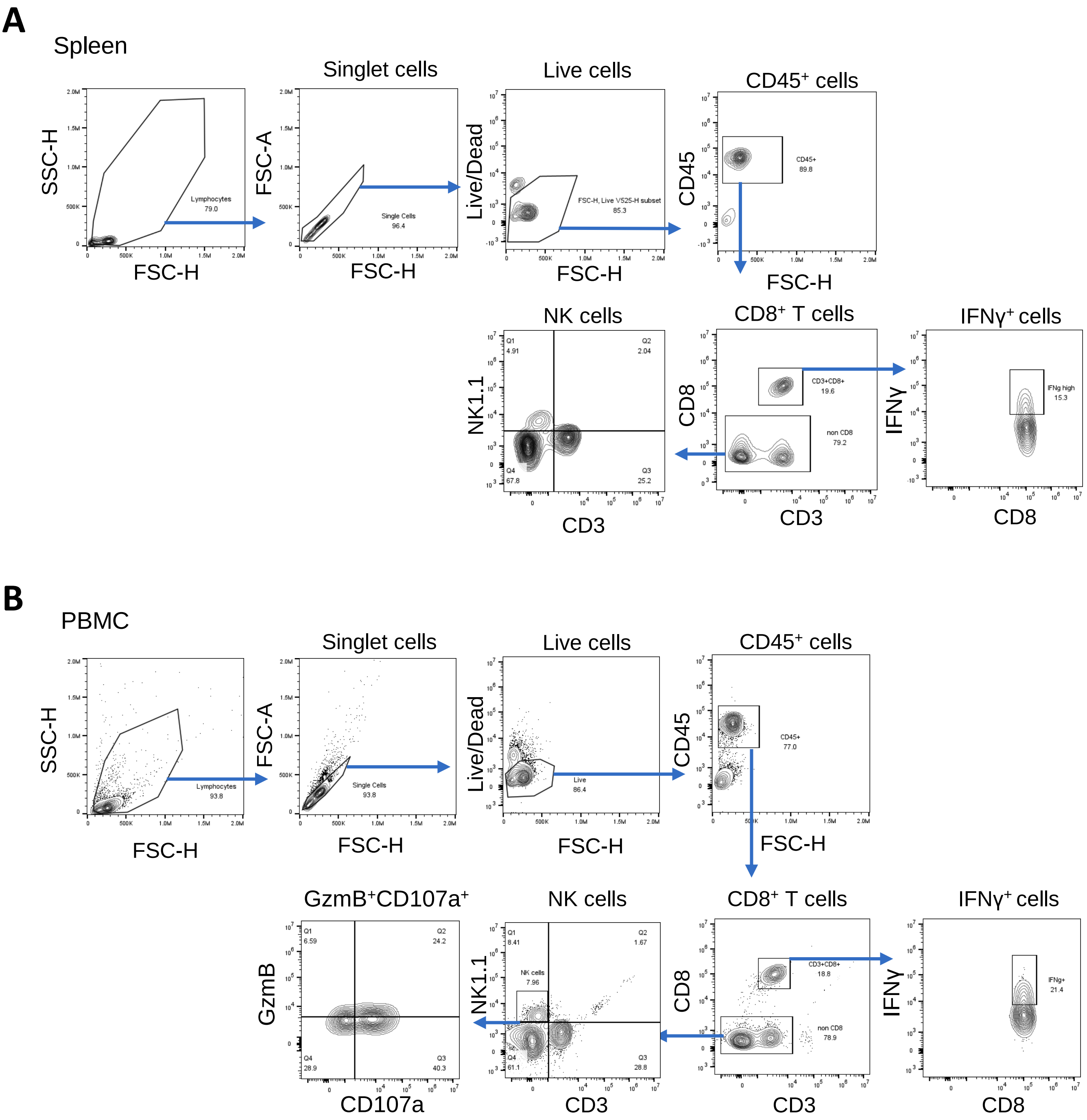


**Supplementary Figure 4. Flow cytometry gating strategy for the characterization of immune cell subsets in spleen cells and PBMCs.** A detailed gating strategy for CD8$^{+}$ T cells and NK cells from spleen cells **(A)** and PBMCs **(B)**, as shown in Figure 5. After live CD45$^{+}$ immune cells gating, each cell subset was gated as follows: CD8$^{+}$ T ; CD3$^{+}$CD8$^{+}$, NK ; CD3$^{-}$ NK1.1$^{+}$
