## Supplementary Table 1 for "Integrating Fas-mediated apoptosis with IFNγ signaling to drive tumor regression in mRNA cancer therapeutics"

Table 1. Primer sequences used in this study.

| | Primer sequence | |
|---|---|---|
| 18S rRNA | F | ACGGAAGGGCACCACCAGGA |
| | R | CACCACCACCCACGGAATCG |
| *Actb* | F | TCACCCACACTGTGCCCATCTACGA |
| | R | CAGCGGAACCGCTCATTGCCAATGG |
| *Ifngr1* | F | TGTGGAGCATAACCGGAGTG |
| | R | GGCTCTTCACAGATCACCGT |
| *Ifngr2* | F | GCCGAACTGTACGGACATCA |
| | R | GCCCAACGGAATCAGGATGA |
| *Irf1* | F | GTGGCAGCCGACACACATCG |
| | R | CTCAGCTGAAGCCCAGGCAGAAA |
| *Cd274* | F | AAAGTCAATGCCCCATACCG |
| | R | TTCTCTTCCCACTCACGGGT |
